# GRASS-NB: Group-structured variable selection for spatial negative binomial data with applications to cancer registry and spatial omics

**DOI:** 10.1101/2025.10.24.684446

**Authors:** Chloe Mattila, Brian Neelon, Kalyani Sonawane, Sha Cao, Peggi Angel, Elizabeth Hill, Souvik Seal

## Abstract

Spatially structured, overdispersed count data with high-dimensional predictors are increasingly observed across studies from population-level epidemiology to cellular-level spatial omics. Feature selection is critical to identify influential predictors, such as key risk factors or biomarkers. Few Bayesian studies have assessed negative binomial regression (NBR) models with standard variable selection priors, like the mixture *spike-and-slab* (SS) or continuous *horseshoe* (HS), but mostly under aspatial settings. Features often form groups; for instance, in population surveys, caloric intake and physical activity may fall under “Diet & Exercise”, while cigarette use and smoking laws belong to “Smoking”. We propose a flexible NBR model that accommodates spatial autocorrelation and introduces a novel group-structured prior by hybridizing SS and HS shrinkage. The model’s performance with different priors is evaluated in terms of specificity, precision, and computational cost under challenging scenarios, including “large *p*, small *n*” cases. We further apply the model to CDC state-level cancer data, comprising demographic, screening, and behavioral covariates, to identify key drivers and population-level risk factors, and to a melanoma spatial omics dataset for predictive modeling expression of gene. An efficient `R` package is provided on `GitHub`.

## 1 Introduction

Datasets with spatially distributed count outcomes and a large number of predictors are routinely collected across diverse scientific domains. In population-level epidemiology, such data arise from large-scale health surveys and cancer registries where outcomes are counts of disease cases or events recorded across geographic regions, such as states or counties (White et al., 2017; Che et al., 2023). In ecology, researchers study species abundance across spatial habitats, often with complex covariates such as climate or land-use measures (Dormann et al., 2007; Michener, 2015). At the cellular scale, spatial omics technologies now produce molecular count measurements (e.g., RNA transcripts) across the tissue or tumor surface (Lewis et al., 2021; Bressan et al., 2023). A common characteristic across these examples is that the outcome is discrete and spatially correlated, often exhibits overdispersion, and the number of predictors (*p*) may be higher than the number of samples (*n*), often with substantial multicollinearity. In such contexts, feature selection, i.e., the process of identifying a subset of predictors that are most relevant to the outcome, becomes crucial for uncovering the most important risk factors in epidemiology or key biomarkers in omics studies (Kirpich et al., 2018; Handorf et al., 2020; Chowdhury and Turin, 2020).

Negative binomial regression (NBR) is a standard approach for modeling overdispersed counts (Hilbe, 2011), extending Poisson regression by introducing a dispersion parameter that accounts for the extra variability. While the NBR model has been extensively used in the above contexts, its integration with high-dimensional variable selection under spatial correlation remains relatively underexplored. From a Bayesian perspective, the variable selection problem is typically addressed through the use of shrinkage priors (Van Erp et al., 2019). Historically, the spike-and-slab (SS) prior (Mitchell and Beauchamp, 1988) has been the most widely used shrinkage prior that assumes each regression coefficient follows a mixture of a point mass at zero (the spike) and a heavy-tailed alternative (the slab). While several novel extensions of this two-part SS prior have been proposed (Ishwaran and Rao, 2005; Malsiner-Walli and Wagner, 2018; Ročková and George, 2018), a class of continuous shrinkage priors has recently emerged as a more computationally efficient alternative. A seminal example is the horseshoe (HS) prior (Carvalho et al., 2009), which provides adaptive shrinkage of coefficients while retaining flexibility for large signals (Bhadra et al., 2019). This prior and its extensions (Piironen and Vehtari, 2017) belong to the class of global-local shrinkage priors (Bhadra et al., 2016), characterized by a “global” hyperparameter that controls overall shrinkage, while “local” hyperparameters control shrinkage per coefficient. In the context of NB regression, a few studies (Dvorzak and Wagner, 2016; Schmidt and Makalic, 2019; Miao et al., 2020; Bhattacharyya et al., 2022) have benchmarked these priors, but only in aspatial settings.

High-dimensional predictors in national survey data, such as state- and county-level demographics, screening, and risk-factor measures from the State Cancer Profiles (SCP) produced by the National Cancer Institute (NCI) and the Centers for Disease Control and Prevention (CDC), typically fall into distinct groups (NCI and CDC, 2025). As an example, the group “Smoking” in the SCP data includes 17 variables, such as the percentages of current smokers, adults currently using e-cigarettes, and states with any 100% smoke-free laws. The variables from the same group tend to be highly collinear, which notoriously complicates variable selection (Segal et al., 2003; Xie and Huang, 2009; Fan and Lv, 2010; Nguyen and Ng, 2020). Standard penalties such as the Lasso can arbitrarily pick one member of a correlated block rather than consistently selecting the full signal set (Zou and Hastie, 2005). To mitigate this, frequentist methods such as the Group Lasso (Yuan and Lin, 2006; Meier et al., 2008; Jacob et al., 2009; Huang et al., 2012) and related shrinkage techniques allow selection at the group level. However, they may struggle when only a subset of members within each group is relevant and provide no insight into the relative importance of individual variables within selected groups. The sparse-group LASSO (Simon et al., 2013; Vincent and Hansen, 2014; Cai et al., 2022) addresses this by combining group- and individual-level penalties in a frequentist framework. However, because the penalties are additive rather than hierarchical, interactions between group- and within-group selection are largely not captured. In the Bayesian literature, truly bi-level shrinkage priors are less common, with a few notable exceptions (Rockova and Lesaffre, 2014; Xu and Ghosh, 2015; Mallick and Yi, 2017).

In the context of bi-level group-structured shrinkage, a hiearchical grouped horseshoe (HS) prior was introduced for variable selection with continuous outcomes (Xu et al., 2016). More recently, Boss et al. (2024) introduced the generalized inverse–gamma–gamma (GIGG) shrinkage prior, which subsumes the grouped HS as a special case under specific parameter choices. The grouped HS has also been adopted in Bayesian neural networks (Ghosh et al., 2019), enabling effective deactivation of individual nodes and entire layers. Building on these ideas, we propose a hybrid prior that uses the horseshoe to control global and group-level importance, and a spike-and-slab component for within-group, variable-level selection. The spike-and-slab layer yields intuitive selection via posterior inclusion probabilities (PIPs), in contrast to reliance on credible intervals in the aforementioned approaches.

Through extensive simulations varying sample size (*n*), number of predictors (*p*), number of groups, and the strength of spatial dependence, we compare ungrouped priors and grouped priors (including our proposed variant) in terms of specificity (exclusion of irrelevant variables), precision (recovery of true signals), and runtime. Applied to CDC State Cancer Profiles (SCP) state-level incidence data for three cancers with grouped predictors (NCI and CDC, 2025), our framework identifies key socioeconomic and screening determinants. We also analyze a cutaneous melanoma spatial transcriptomics (ST) dataset (Thrane et al., 2018), predicting the expression of pathology-guided region-specific marker genes from biologically relevant predictor sets. To our knowledge, we provide the first R package for Bayesian variable selection in spatially indexed count data—GRASS-NB—which offers flexible prior specifications.

## 2 Methods

### 2.1 Model Specification

Suppose there are *n* locations. For each location *i*, we observe a count *y*_*i*_ and a *p*-dimensional feature vector **K**_*i*_ with first element 1, for *i* = 1,…, *n*. We assume *y*_*i*_ follows a negative binomial (NB) distribution with failure probability *ψ*_*i*_ ∈ [0, 1] and overdispersion parameter *r >* 0. To facilitate Bayesian computation using the Pólya-Gamma data augmentation scheme (Pillow and Scott, 2012), we model *ψ*_*i*_ with an inverse-logit link,

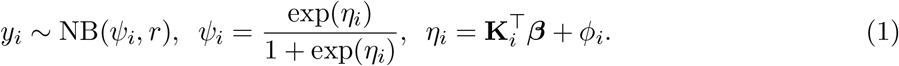

The coefficient vector is ***β*** = (*β*_0_,…, *β*_*p*−1_)^*T*^, where *β*_0_ is the intercept and the remaining coefficients 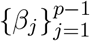 correspond to the feature effects. *ϕ*_*i*_ denotes a spatial random effect, which varies smoothly along the neighborhood of each location. We place an intrinsic conditional autoregressive (ICAR) prior (Banerjee et al., 2014) on 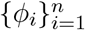:

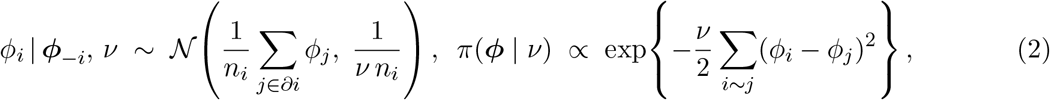

where ***ϕ*** = (*ϕ*_1_,…, *ϕ*_*n*_)^⊤^ and 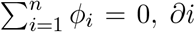 is the neighbor set of node *i, n*_*i*_ = |*∂i*|, and *i ∼ j* denotes adjacency on the (unweighted) neighborhood graph. Equivalently, the precision matrix is *ν***Q** with **Q** = *D* − *W*, *W* is the adjacency matrix following the queen contiguity where two regions are considered neighbors if they share a border or vertex, which reflects the neighborhood structure of the contiguous United States. *D* is a diagonal matrix of neighbor counts; *W*_*ij*_ = 1 if *j* ∈ *∂i* or 0 otherwise, and 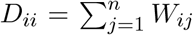. Since the ICAR likelihood is improper, we add a small “nugget” precision (*ϵ* = 10^−8^) to the diagonal of the precision matrix (Banerjee et al., 2014). The constraint 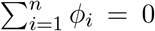 separates the spatial effects from the intercept *β*_0_ and ensures identifiability. The precision parameter *ν >* 0 controls the overall degree of spatial smoothing, with smaller *ν* enforcing stronger similarity across neighboring areas and larger *ν* allowing greater spatial heterogeneity. Next, we discuss the variable selection priors for *β*.

### 2.2 Variable Selection Priors

#### 2.2.1 Standard Priors

First, we consider the standard horseshoe (HS) (Carvalho et al., 2009) and spike-and-slab (SS) (Ishwaran and Rao, 2005) priors:

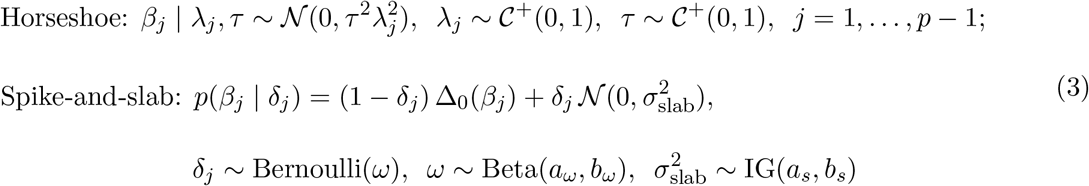

*τ* ^2^ is the global hyperparameter, controlling the overall shrinkage, reflecting prior belief about coefficient sparsity. 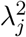 is the local hyperparameter that counteracts *τ* ^2^ for truly large effects, allowing them to escape global shrinkage. In the SS prior, each coefficient is governed by an inclusion indicator: *δ*_*j*_ *∼* Bernoulli(*ω*), where *ω* the prior fraction of active coefficients. Δ_0_(.) is the Dirac delta function, which places all its mass at zero, enforcing sparsity by shrinking small effects exactly to zero. The slab, 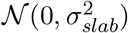, allows flexibility of the active features over a broad Gaussian prior of plausible coefficients.

#### 2.2.2 Group Structured Priors

Suppose the features belong to *G* known groups (see Fig. 4). A group *g* contains *M*_*g*_ features, and 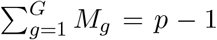. We introduce two structured priors accounting for the group hierarchy of the features,

Grouped horseshoe: 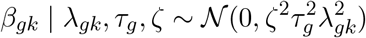,

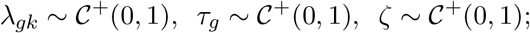

Grouped spike-and-slab (proposed): 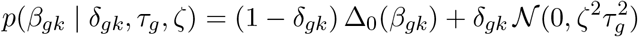,

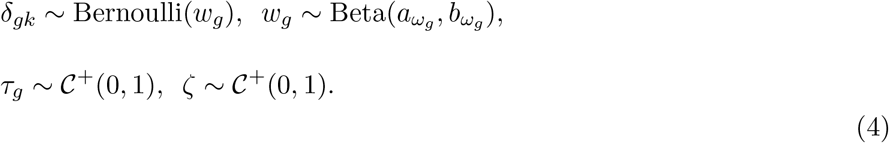

For grouped horseshoe (HS) (Xu et al., 2016), the “global-global” hyperparameter, *ζ*, controls overall shrinkage. 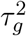 is a group-specific “global-local” hyperparameter determining the relevance of group *g* as a whole. Finally, 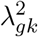 is a within-group “local-local” hyperparameter, which allows individual coefficients within active groups to escape shrinkage. This multilevel extension enables flexible shrinkage at the global, group, and feature levels, accommodating the hierarchical structure common in many datasets. However, variable selection under this prior—and under its GIGG extension (Boss et al., 2024)—is typically conducted via post hoc credible-interval analysis, a procedure that is inherently ad hoc and sensitive to the specified interval width. However, variable selection under this prior—and under its GIGG extension (Boss et al., 2024)—is typically conducted via post hoc credible-interval (CrI) analysis, a procedure that is inherently ad hoc and sensitive to the specified interval width.

To address this limitation, we propose a hybrid group-structured prior that integrates the adaptive shrinkage of the horseshoe (HS) with the interpretability of the spike-and-slab (SS) framework. Specifically, we employ a group-level HS structure to regulate overall shrinkage and group importance, while embedding an SS component within each group to enable variable-level selection through posterior inclusion probabilities (PIPs). More concretely, the slab variance is factorized as 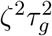, where 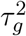 captures the relevance of group *g* and *ζ*^2^ controls global shrinkage, mirroring the grouped HS construction. Variable inclusion is determined based on a PIP threshold (e.g., 0.5 for the median probability model), providing a more principled and transparent alternative to CrI-based selection. As shown later, the proposed grouped SS prior also achieves modest improvements in power over the grouped HS prior. For singleton groups, we drop the redundant level: in the grouped HS, we fix the local scale to one (*λ*_*gk*_ = 1); in the grouped SS, we fix the group scale to one 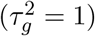. We derive a Gibbs sampler with closed-form full conditional distributions for all parameters. Convergence is assessed using trace plots and the Geweke statistic (Geweke, 1992), implemented in the `coda` package (Plummer et al., 2006).

## 3 Results

### 3.1 Simulation Study

The first simulation evaluated the impact of modeling versus ignoring spatial dependence under a true spatial random effect with multicollinear features and no known group structure. In the second simulation, we imposed a hierarchical feature structure and compared model performance under group-structured priors versus standard priors. The final simulation introduced singletons groups alongside a hierarchical feature structure and a true spatial random effect, allowing comparison of group-structured and standard priors under spatial dependence.

#### 3.1.1 Impact of Explicitly Modeling Spatial Dependence

We generated synthetic datasets with sample sizes *N* ∈ {50, 100} and feature dimensions *p* ∈ {50, 200}. Locations were placed on a 10 *×* 5 grid for *N* = 50 and on a 10 *×* 10 grid for *N* = 100. The design matrix **K** consisted of (*p*− 1) continuous features, and a column of all 1’s (for intercept). The features were simulated from 𝒩_*p*−1_(**0**, 0.1 **Σ**), where Σ_*jk*_ = *ρ*^|*j*−*k*|^ for *j, k* = 1,…, *p* − 1 (so Σ_*jj*_ = 1). We varied *ρ* ∈ {0, 0.25, 0.5} to induce varying degrees of multicollinearity. Note that there is no explicit group structure between the features. We randomly selected ten features to have non-null effects and fixed their coefficients at *β*_*j*_ = 2. The spatial random effects (SRE) vector,***ϕ*** = (*ϕ*_1_,…, *ϕ*_*n*_)^⊤^, was simulated from an ICAR prior, ***ϕ*** | *ν ∼* 𝒩_*n*_ **0**, [*ν*(**Q** + *ϵ***I**_*n*_)]^−1^, as derived from equation 2 with a nugget *ϵ* = 10^−8^. **Q** was constructed from a queen-contiguity adjacency. We considered *ν* ∈ {0.1, 0.25}, reflecting stronger and weaker spatial dependence, respectively. Finally, the count outcome *y*_*i*_ was sampled from the NB model in equation 1 with *r* = 1.

We fitted four models: (i) two with the standard HS and SS priors from Eq. 3 on *β*_*j*_’s *with* the SRE, and (ii) two using the same priors *without* the SRE. In addition, we used the `R package pogit` (Dvorzak and Wagner, 2016), which is the only publicly available tool offering Bayesian variable selection for count data. Specifically, we used their function `negbinBvs`, which implements the standard SS prior but does not accommodate spatial effects. We evaluated performance using receiver operating characteristic (ROC) curves and area under the curve (AUC). For the SS model, decision thresholds were given by posterior inclusion probabilities 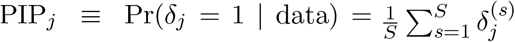, where *S* denotes the post-burn-in iterations. For the HS model, thresholds were based on the widths of equal-tailed-interval (ETI) credible intervals (Makowski et al., 2019). Results are based on 10,000 MCMC iterations with a 50% burn-in. HS consistently ran faster than SS. At *N* = 100 and *p* = 50, HS without SRE ran in a median of 43.0 seconds versus 118.7 seconds for SS. As dimensionality rose to *p* = 200, HS required 54.2 seconds, while SS increased sharply to 374.9 seconds (7 times), underscoring HS’s superior scalability. Analyses were performed on a workstation running Microsoft Windows 11 with an Intel® Core™ i7-14700 processor (20 cores, 2.10 GHz) and 32 GB RAM, using R version 4.5.1.

Fig. 1 shows that models with the SRE (solid lines) achieved higher selection accuracy than those without (dashed lines). The gap widened as feature correlation (*ρ*) increased, indicating that modeling residual spatial dependence is essential, particularly in highly multicollinear settings. As expected, all models performed better under lower spatial dependence (*ν* = 0.25). The `pogit` SS implementation was marginally different than our SS model without the SRE, possibly due to differences in data augmentation strategies (finite mixtures (Dvorzak and Wagner, 2016) versus Pólya-Gamma).

**Figure 1.**
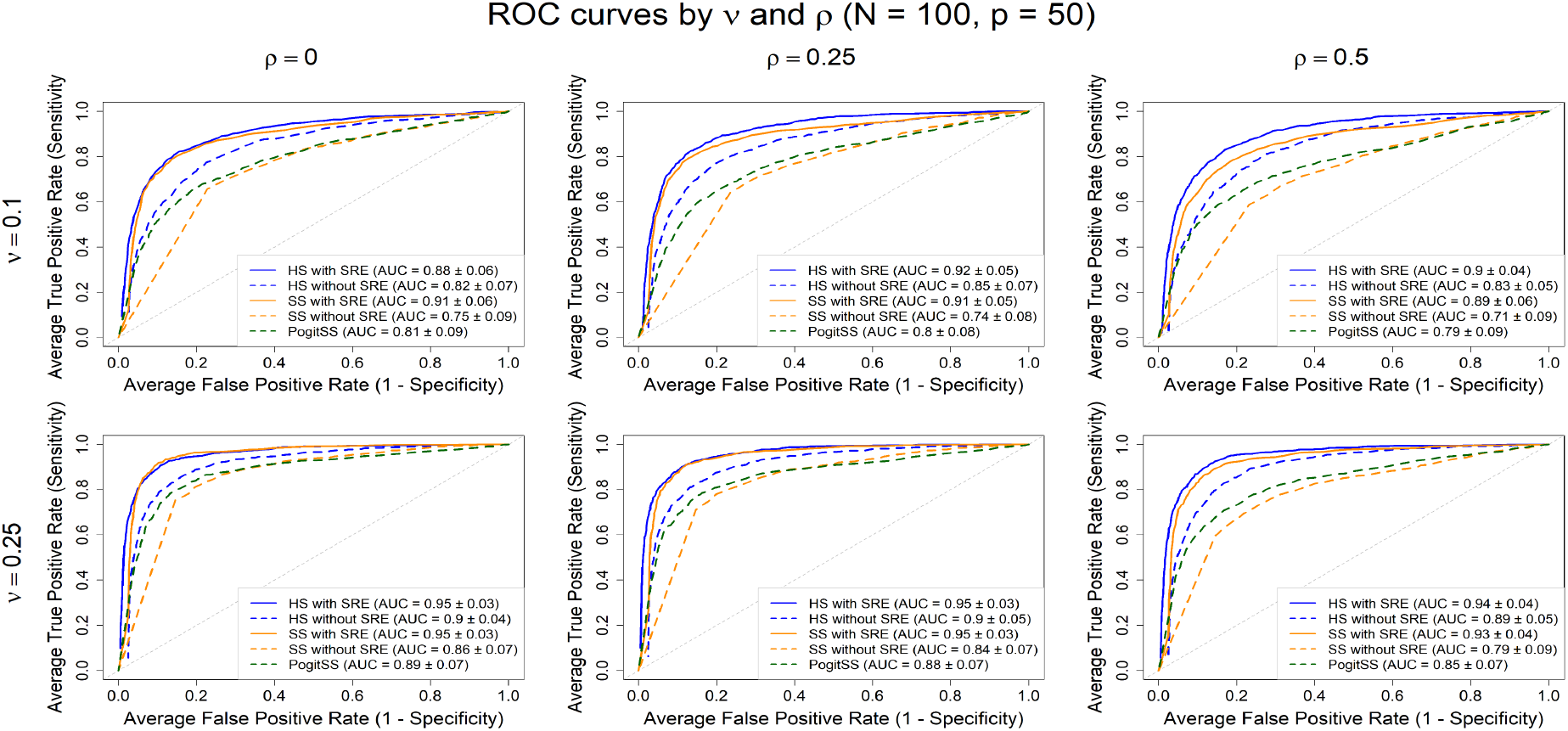
ROC curves of the models under high (top row) and low (bottom row) spatial dependence. HS/SS with SRE outperform no-SRE models, especially under high spatial dependence.

#### 3.1.2 Impact of Using Structured Priors

We partitioned the features into *G* groups (Section 2.2.2). Group *g* contains *M*_*g*_ features that are correlated within the group, whereas distinct groups are mutually independent. The feature matrix **K** was partitioned as **K** = [**1** | **K**_1_ |…| **K**_*G*_]. For each group *g*, the *i*th row of the *n × M*_*g*_ submatrix **K**_*g*_ was simulated with an AR(1) correlation structure: 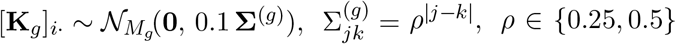. We set the group size to *M*_*g*_ = 5, yielding *G* = ⌈(*p*)*/M*_*g*_⌉ groups, with all groups of size five except one, which had four. We considered two signal configurations (10 non-null coefficients in total): (1) two randomly selected groups *fully* active, and (2) three randomly selected groups *partially* active, with some within-group coefficients set to zero. For each non-null coefficient *β*_*j*_, the signal magnitude was independently drawn from Unif(0.5, 2); all remaining coefficients, intercept included, were set to zero.

We fit four models with different prior structures: HS and SS priors *with* grouping (Eq. 4) and the standard HS and SS priors *without* grouping (Eq. 3). From Fig. 2, in both cases, grouped priors achieved better performance than their non-grouped counterparts. Gains were largest in Case 1, where two entire groups were active. Performance improved with larger *N*, as expected, while predictor correlation had a negligible impact. Overall, the results suggest that explicitly modeling feature–group structure improves selection performance, especially when *p > n*, by borrowing strength within groups to identify active features and “switch off” entirely inactive groups. In select cases, the grouped SS prior marginally outperformed the grouped HS.

**Figure 2.**
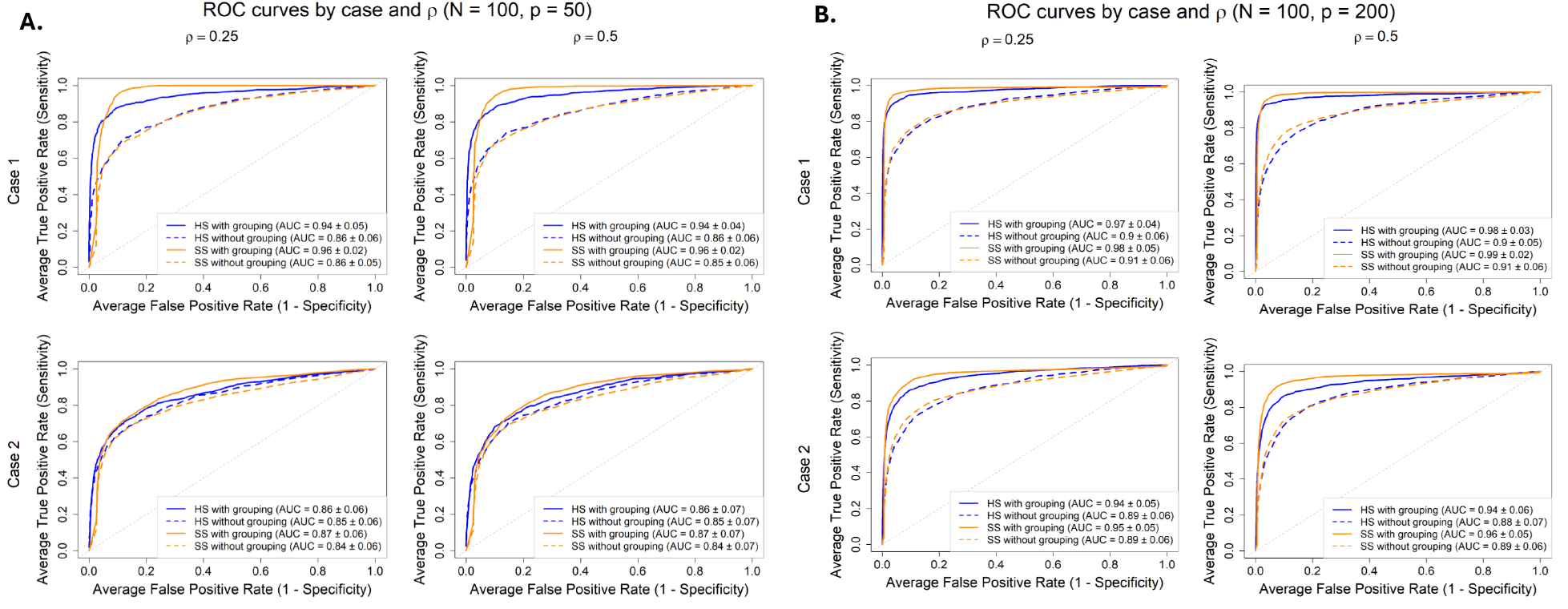
ROC curves for (A) *p* = 50, (B) *p* = 100 with 10 non-nulls total. Case 1 (top row), two full groups active; Case 2 (bottom row), three groups partially active. HS and SS with grouping outperform methods without grouping, especially for large *p*.

#### 3.1.3 Impact of Spatial Modeling and Structured Priors with Singleton Groups

We considered *p* − 1 non-intercept features. The first *p/*2 features were partitioned into groups of equal size (*M*_*g*_ = 5), with features within the same group exhibiting an intra-group correlation of *ρ* = 0.5 and distinct groups treated as mutually independent, as in Section 3.1.2. The remaining (*p/*2) − 1 features were treated as singletons, each forming its own group, and were generated from a correlated structure with pairwise correlations *ρ* = 0.5 among all singleton features, following Section 3.1.1. A total of ten non-null coefficients were included, split evenly between the grouped and singleton features at random. For the grouped features, the two signal configurations considered in Section 2.2.2 were again considered. Each non-null coefficient *β*_*j*_, were fixed at four (*β* = 4).

As in Section 3.1.2, HS and SS priors *with* grouping (Eq. 4) and the standard priors *without* grouping (Eq. 3) were fit with spatial random effects (SRE). Figure 3, displays the ROC curves for *N* = 100 under the two signal configurations for the *p/*2 grouped features at *p* = 50 (left) and *p* = 200 (right). In Case 1, the SS model with grouping and SRE achieved the highest AUC across both dimensions, indicating that explicitly modeling group structure was beneficial when all features within a group were truly non-null. In Case 2, however, the SRE-only models slightly outperformed the grouping and SRE models, suggesting that group-level shrinkage can be detrimental when only a subset of features within a group is active and the overall signal is sparse. In this scenario, partial activation within a group effectively dampened the signal, reducing model discrimination. Across both signal configurations, the SS prior generally produced slightly higher or comparable AUCs than the HS prior, particularly in higher-dimensional settings (*p* = 200). This simulation represented a particularly challenging scenario, combining spatial dependence, correlated singleton features and partial group activation, so the moderate AUC values observed here in Figure 3 are expected and reflect the inherent difficultly of distinguishing weak and diffuse signals under such complex correlation structures.

**Figure 3.**
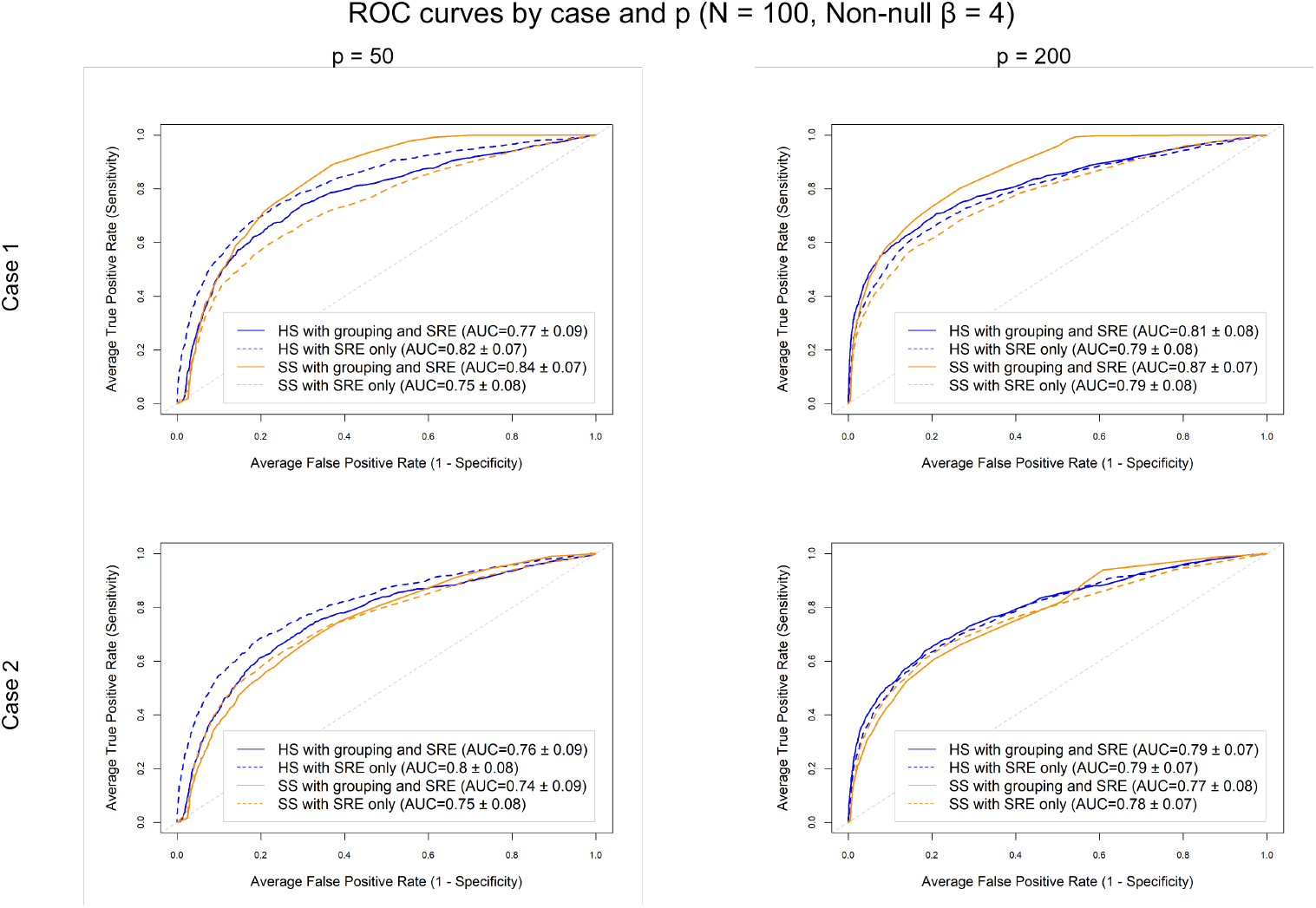
ROC curves for *p* = 50 (left) and *p* = 200 (right) for *N* = 100 and ten non-nulls (effect size = 4), split evenly between the grouped features and singleton features. Case 1 (top row), one fully active group; Case 2 (bottom row), 2 groups partially active. In Case 1, the SS model with grouping and spatial effects achieved the highest AUCs, indicating that leveraging group structure improves detection when signals are concentrated within groups. In Case 2, however, the spatial-only models perform better, suggesting that group-level shrinkage can hinder detection when signals are sparse and correlated singletons are present.

**Figure 4.**
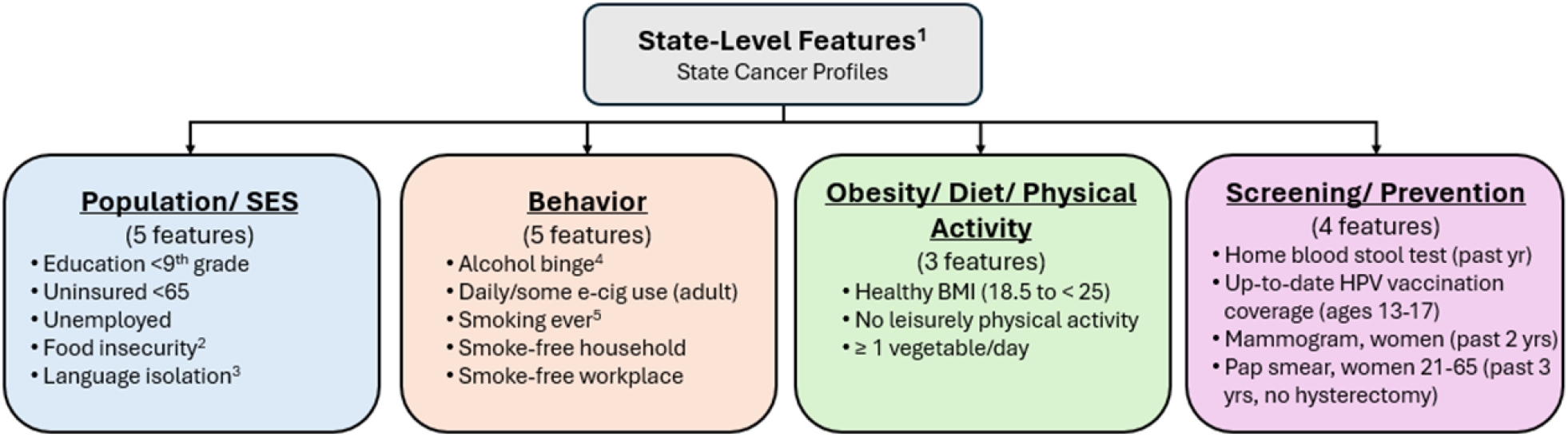
Conceptual groupings of 17 state-level features. ^1^Proportions. ^2^Limited food access: uncertainty in adequate, safe food. ^3^Language isolation: no one (14+) speaks only English or speaks it well (household). ^4^Alcohol binge: ≥4 drinks (women) or ≥5 drinks (men) on one occasion. ^5^Smoking ever: ≥100 cigarettes in a lifetime.

### 3.2 Real Data Application

#### 3.2.1 State-level cancer incidence data

We analyzed state-level incidence counts among U.S. adults aged ≥ 50 years for three common cancers, female breast, prostate, and bladder, that together account for a substantial share of the cancer burden (McMillan et al., 2025). Counts were obtained from the SCP website (NCI and CDC, 2025), as five-year average (2017–2021). Figure 6A displays incidence rates per 100,000 by state, i.e., 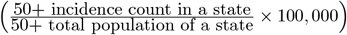. Analyses were restricted to the contiguous U.S., excluding Alaska and Hawaii, and Indiana (data unavailable). State-level features were also collected from the same website, spanning 2019-2023. Many of the 30 initial features were nearly collinear (pairwise correlation ≈ 1). We reduced redundancy using a correlation dendrogram (Gu et al., 2016), retaining 17 features with pairwise correlations below 0.75. Based on the SCP portal and our interpretation, features were organized into four nominal groups: Population/SES, Behavior, Obesity/Diet/Physical Activity, and Screening/Prevention (Fig. 4). All features, originally expressed as percentages, were log-transformed and standardized (mean = 0, variance = 1). State-level counts of adults aged 50+ were obtained from the U.S. Census Bureau’s 2020 *State Population by Characteristics* dataset (U.S. Census Bureau, 2025), which is based on the 2020 Census and accounts for births, deaths, and migration.

We fit the NBR model from Eq. 1, additionally offsetting for the total population of 50+ adults in each state (*P*_*i*_) and scaling outcomes to rates per 100, 000:

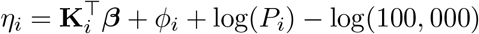

where, for state 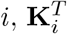 is a vector of state-level features (grouped as in Fig. 4) plus a fixed intercept, *ϕ*_*i*_ is the state-specific spatial random effect. For methodological comparability, we used a common feature set across all cancers; however, sex-specific variables (e.g., “Mammogram, past 2 yrs”) are unlikely to be directly relevant for cancers in the opposite sex, though they may still serve as proxies for broader group-level factors (e.g., screening and prevention practices). We considered the grouped HS and SS priors from Eq. 4, and 50, 000 MCMC iterations (50% burn-in). Trace plots and Geweke statistics indicated convergence for all features. The posterior mean estimates from the HS and SS models largely coincided (Fig. 6D) and the estimated *ϕ*_*i*_’s were likewise consistent (Fig. 6B, C). For inference, we used 95% credible intervals for HS and a PIP threshold of 0.6 for SS.

For breast cancer (light pink in Fig. 5), all 95% credible intervals under the HS model contained zero. At the PIP threshold of 0.6, the SS model indicated a negative association (*β*_*j*_ *<* 0) between “Smoking banned at work” and breast cancer incidence rates, and a positive association (*β*_*j*_ *>* 0) with “Pap smear past 3y (21-65)”. States with workplace smoking bans tend to exhibit lower expected breast cancer incidence among women 50+, consistent with broader public health and behavioral environments favoring reduced cancer risk. In contrast, higher pap smear screening rates were associated with a greater recorded incidence, likely reflecting enhanced healthcare access and detection rather than increased biological risk.

**Figure 5.**
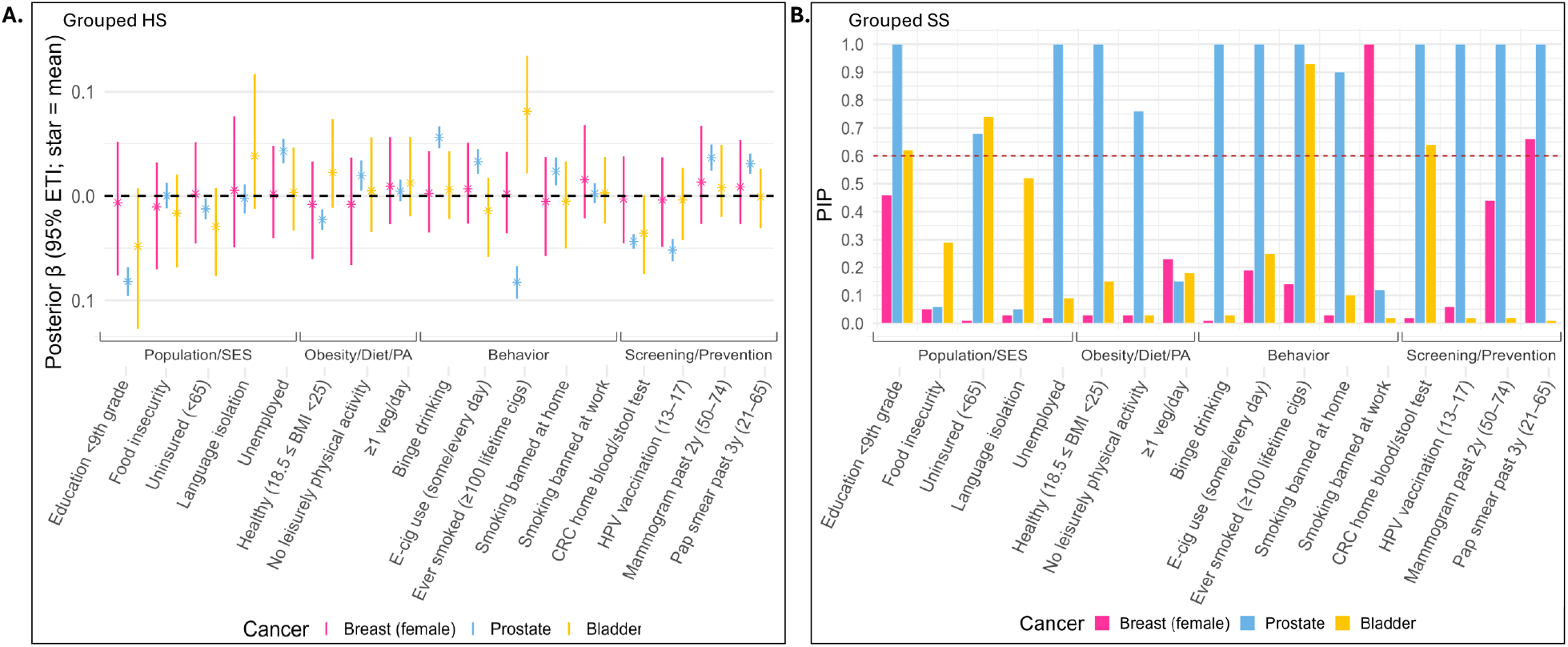
(A) Grouped HS 95% credible intervals (CrI) for features in the cancer application. (B) Grouped SS posterior inclusion probabilities (PIP), red dashed line marks the 0.6 threshold.

For prostate cancer (light blue), thirteen HS-based 95% credible intervals excluded zero. Notably, several of the negative associations, in Fig. 5, correspond to lower education, reduced screening, and lower HPV vaccination coverage, whereas positive associations involve higher unemployment and binge drinking. These patterns suggest that limited preventive care and screening in states with lower education and vaccination rates contribute to observed prostate cancer incidence, while socioeco-nomic stressors and high-risk behaviors are linked to adverse health outcomes. Although the negative association with smoking appears counterintuitive, it may reflect a competing-risks/detection mechanism: higher smoking-related mortality reduces the opportunity for a prostate cancer diagnosis, yielding lower recorded incidence. At a PIP threshold of 0.6, the SS model selected the same features as the grouped HS model. Notably, three Population/SES variables, two Obesity/Diet/Physical Activity variables, four Behavioral variables, and all four Screening/Prevention variables had PIP *>* 0.6, underscoring the overall importance of these groups for prostate cancer incidence (Coughlin, 2020; Jain et al., 2023).

For bladder cancer (yellow), the HS model selected only “Ever smoked (≥ 100 lifetime cigarettes),” whereas the SS model selected four features across 3 of the conceptual groupings (Table 1. While the selected features reflect the overall influence of the Population/SES group (Sung et al., 2019) and smoking is a well-established risk factor (Freedman et al., 2011), the stool–blood testing variable likely serves only as a proxy for screening practices or healthcare access. Overall, although posterior mean estimates from the HS and SS models largely coincided, the grouped SS appeared more powerful across the three cancers.

**Table 1.** Selected features, shown in Fig. 5, by cancer and variable-selection method. For grouped HS, selected features have 95% CrI excluding zero; for grouped SS, selected features have PIP ≥ 0.6. Color-coded boxes indicate conceptual feature groups consistent with Fig. 4 (blue: Population/SES; green: Obesity/Diet/PA; orange: Behavior; purple: Screening/Prevention).

| Cancer Type | Grouped Horseshoe | Grouped Spike-and-Slab |
| --- | --- | --- |
| Breast (female) | — | Smoking banned at work |
|  |  | Pap smear past 3y (21–65) |
| Prostate | Education < 9th grade,<br>Uninsured (< 65), Unemployed | Education < 9th grade,<br>Uninsured (< 65), Unemployed |
| | Healthy ($15.5 \leq \text{BMI} < 25$),<br>No leisurely physical activity | Healthy ($15.5 \leq \text{BMI} < 25$),<br>No leisurely physical activity |
| | Binge drinking,<br>E-cig use (some/every day),<br>Ever smoked ($\geq 100$ lifetime cigs),<br>Smoking banned at home | Binge drinking,<br>E-cig use (some/every day),<br>Ever smoked ($\geq 100$ lifetime cigs),<br>Smoking banned at home |
|  | CRC home blood/stool test, HPV<br>vaccination (13–17),<br>Mammogram past 2y (50–74), Pap<br>smear past 3y (21–65) | CRC home blood/stool test, HPV<br>vaccination (13–17),<br>Mammogram past 2y (50–74), Pap<br>smear past 3y (21–65) |
| Bladder | Ever smoked ($\geq 100$ lifetime cigs) | Education < 9th grade,<br>Uninsured (< 65) |
| | | Ever smoked ($\geq 100$ lifetime cigs) |
|  |  | CRC home blood/stool test |

#### 3.2.2 Spatial transcriptomics data

We analyzed a cutaneous melanoma dataset (Thrane et al., 2018) generated with spatial transcriptomics (ST) (Ståhl et al., 2016) from a long-term survivor (*>* 10 years). The dataset comprises 293 spots (100 *µ*m spot diameter; 200 *µ*m center-to-center spacing) and 16,148 genes. After quality control, removing genes expressed in fewer than 20% of spots, we retained 5,296 genes. Using a Poisson factor analysis model (Berglund et al., 2018), Thrane et al. (2018) identified three prominent spatial factors (Fig. 7A), corresponding to three regions pathologically annotated as “Melanoma-A,” “Transition area,” and “Lymphoid.” Each factor was associated with enriched genes (Fig. 7A); in this analysis, we focus on the top two genes for each factor: CD63 and PMEL for Melanoma-A, FTL and B2M for Transition area, and ACTB and CD74 for Lymphoid (Fig. 7C). Notably, the expression patterns of these genes can be interpreted as proxies for the three regions. Our goal was to determine whether these genes can be reliably predicted by a subset of genes (Zhou et al., 2025). To define a biologically grounded candidate set, we selected genes annotated to Gene Ontology (GO) terms reflecting core melanocytic programs (used keywords: “melanin,” “melanosome,” “pigment,” “melanocyte,” “melanocyte differentiation”). These terms summarize the lineage-specific processes underlying pigment production and organelle biogenesis in melanoma cells, and led to 23 GO terms with 74 genes (Fig. 7B). Because our approach does not allow overlap between predictor groups, any gene annotated to multiple GO terms was randomly assigned to a single term. We fit the models using the proposed grouped SS prior and declared variables significant at a PIP threshold of 0.6, as earlier.

**Figure 6.**
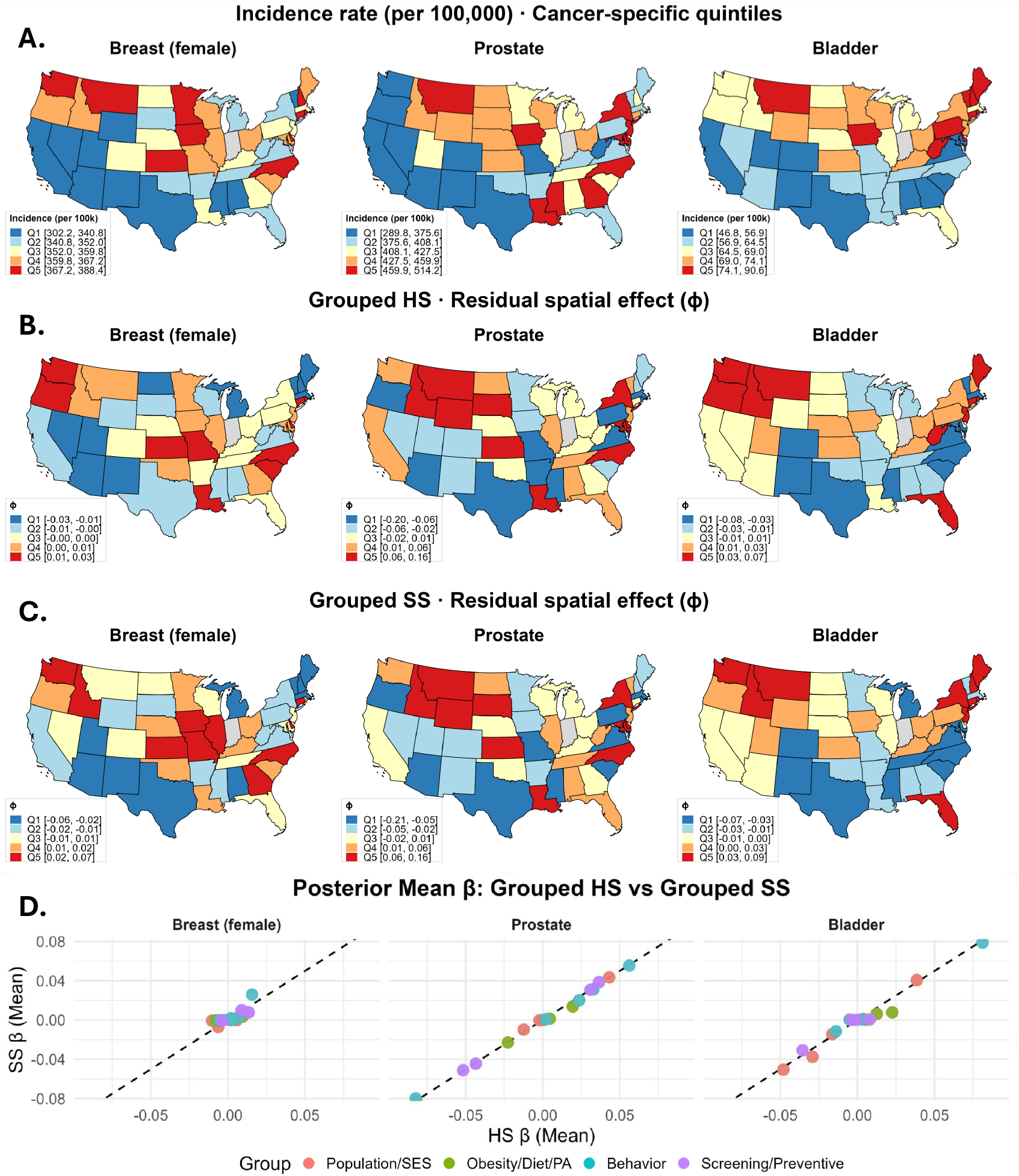
(A) State-level cancer incidence rates (per 100,000) among adults aged 50+ in the contiguous U.S., for breast (female), prostate, colorectal, and bladder cancers. Rates are 5-year averages (2017–2021) from the State Cancer Profiles; darker red indicates higher incidence (quintiles). Indiana is shaded gray where data are unavailable. (B) Posterior mean spatial effects (*ϕ*) from the grouped Horseshoe (HS) model. (C) Posterior mean spatial effects (*ϕ*) from the grouped Spike-and-Slab (SS) model. Legends show quintiles, with darker red indicating stronger positive spatial effects. (D) Scatterplots of posterior mean regression coefficients comparing HS and SS. Each point is a predictor, colored by conceptual group. The dashed line indicates perfect agreement (HS = SS).

**Figure 7.**
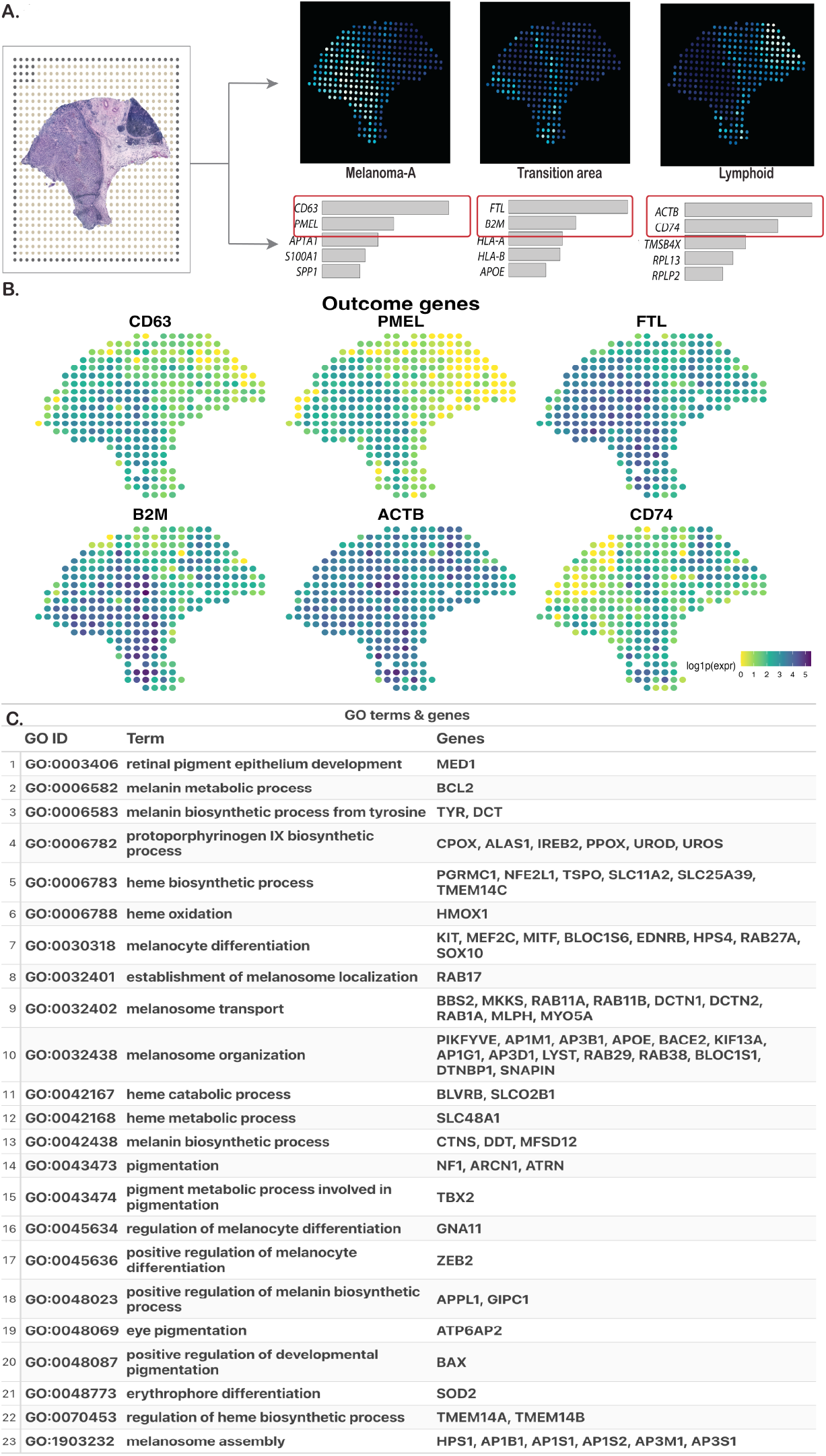
(A) Histology image and three spatial factors, with two marker genes highlighted (from Thrane et al. (2018)). (B) Predictor GO terms and corresponding genes. (C) Spot-level log(expression + 1) profiles for the marker genes.

For CD63 and PMEL, the GO term `GO:0006583` (melanin biosynthetic process) and its associated genes *TYR* and *DCT* were positively associated (Figs. 8A and B), suggesting activation of the tyrosine-to-melanin conversion pathway in the Melanoma-A region. A quick look at *DCT* expression (Fig. 8B) confirms that it is indeed overexpressed in that region. For CD63 specifically, three genes annotated to `GO:0032438` (melanosome organization) were significant, which is unsurprising as this term regulates structural and trafficking processes essential for melanosome assembly. From Fig. 8D, the residual spatial surface (scaled by its standard deviation) aligns with the Melanoma-A region. Moreover, the mean absolute residual, 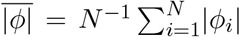, is larger for PMEL than for CD63 (0.28 vs. 0.12), indicating poorer predictive performance (i.e., larger unexplained spatial signal) for PMEL. For FTL and B2M, `GO:0032438` was again significant (6 genes for FTL and for B2M), indicating activation of the pathway also in the Transition area. `GO:0048069` (eye pigmentation; gene *ATP6AP2*) and `GO:0048773` (erythrophore differentiation; gene *SOD2*) were also significant for B2M. *ATP6AP2* (the prorenin receptor) has emerged as a biomarker in multiple cancers, including melanoma (Wang et al., 2020), and *SOD2* has been reported to play context-dependent roles in melanoma, with studies supporting both tumor-suppressive and pro-tumorigenic effects (Carvalho et al., 2022). The residual spatial surface for B2M aligned well with the Transition area, whereas the alignment was less pronounced for FTL. For ACTB and CD74, *APOE* from `GO:0032438` (melanosome organization) was significant (with two additional `GO:0032438` genes for ACTB). *MEF2C*, a MADS-box transcription factor (Lin et al., 1998), from `GO:0030318` (melanocyte differentiation) was significant for both, which is plausible given the heightened transcriptional activity in the Lymphoid region. Interestingly, the only two significant negative associations were observed for CD74, namely with *DCT* and *SOX10*. This is consistent with their expression patterns: both genes are enriched in the Melanoma-A region and largely absent from the Lymphoid (Fig. 8B). The residual spatial surface for CD74 aligned with the Lymphoid region, whereas ACTB did not, consistent with its near-uniform expression across the tissue (Fig. 7C). Overall, these results illustrate how principled spatial modeling, coupled with our group-structured shrinkage prior, can uncover meaningful predictive signals in ST datasets.

**Figure 8.**
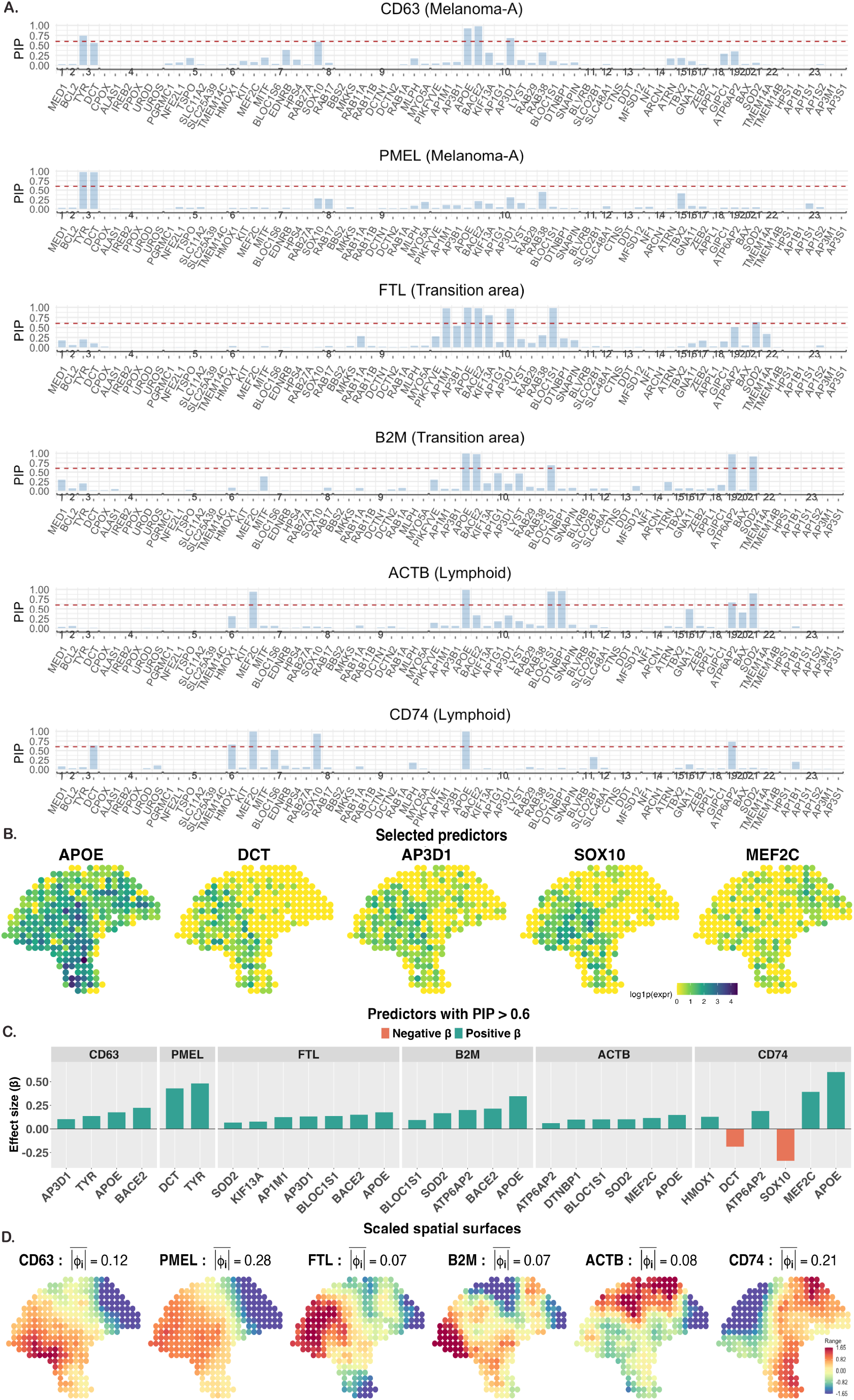
(A) PIP results for the marker genes as outcomes, with the red-dotted line denoting a probability of 0.6. (B) Expression profile of select predictors. (C) Effect sizes of select significant predictors. (D) Scaled residual spatial surface of each outcome gene.

## 4 Discussion

In this study, we investigated variable selection for spatially indexed negative binomial outcomes when predictors are naturally organized into correlated groups—a setting common in national cancer registry surveillance and spatial omics. We compared two widely used ungrouped priors, the continuous-shrinkage horseshoe (HS) and the discrete-mixture spike-and-slab (SS), as well as a grouped HS prior (Xu et al., 2016), and introduced a hybrid, group-structured prior that fuses the strengths of HS and SS. By explicitly encoding known feature–group structure, the proposed prior, together with the negative binomial regression framework, offers a practical and interpretable approach for high-dimensional spatial count data.

Our simulation results indicate thatexplicitly modeling spatial dependence improves variable-selection performance for both priors when a count outcome is spatially distributed. When variables form groups with strong within-group but negligible between-group correlation, encoding this structure, as in the grouped HS and proposed grouped SS priors, substantially improves signal recovery, particularly when *p > n*. In the SCP cancer data for breast (female), prostate, and bladder cancers, state-level incidence exhibited pronounced spatial patterns across the United States, necessitating explicit spatial modeling. Across the four groups, Population/SES, Obesity/Diet/Physical Activity, Behavior, and Screening/Prevention, totaling 17 features, both the grouped HS prior (based on 95% credible intervals) and the proposed grouped SS prior (based on posterior inclusion probability (PIP) *>* 0.6) identified features from the Behavior group to be the most influential predictors of state-level breast (female), prostate, and bladder cancer incidence, though grouped HS selected no features for breast (female). However, the grouped SS consistently selected as many or more features across all cancer types, additionally identifying the Screening/ Prevention group as important for all cancers. These results suggest that the grouped SS prior offers improved power in identifying relevant features while maintaining interpretability through PIPs. Notably, strong between-group correlations made the analysis substantially more challenging than the near-independent group settings for which our extensions are well-suited. A natural remedy could be to define groups via correlation-based clustering rather than nominal categories, though this may reduce interpretability. In the spatial transcriptomics application, our proposed prior reliably recovered biologically mean-ingful predictor genes from large gene ontology (GO)-derived candidate sets. This paves the way for concise, interpretable models, with uncertainty quantification, that preserve specificity while scaling to high-dimensional gene panels.

The grouped HS and SS priors performed similarly in our experiments, with SS showing a modest advantage in some settings (e.g., select simulations and the SCP analysis), elevating computational efficiency as a key consideration. The HS prior admits a highly efficient Gibbs sampler that runs in a fraction of the SS runtime, making HS a sensible default when *p* is large (e.g., *p* ≥ 200). By contrast, the SS prior offers more direct interpretability via PIPs. In future work, we will pursue variational inference for the grouped SS prior following Miao et al. (2020). More importantly, we aim to incorporate between-group feature correlations more directly into our framework. Motivated by Ma et al. (2024), we will place a correlation-aware prior on the SS inclusion indicator vector ***δ***. Another important direction for future work is to incorporate spatially varying coefficients (Gelfand et al., 2003) within the NB regression framework, augmenting the proposed prior, following Seal and Neelon (2025). We release an open-source `R package GRASS-NB on GitHub`, which implements (i) the standard horseshoe (HS) and spike-and-slab (SS) priors, (ii) grouped HS, and (iii) proposed grouped SS, each with a spatial random effect, in a negative binomial regression model.

## 5 Data and software availability

A GitHub *R* package named `GRASS-NB`, with the proposed method and the first two datasets in “.rda” format, is available at https://github.com/Cmattila/GRASS-NB/. The original melanoma dataset is available at the link: https://zenodo.org/records/8215682 with sample ID “`ST_mel1_rep2`“.

## 6 Funding

C.M., B.N., E.H., and S.S. were supported in part by the Biostatistics Shared Resource, Hollings Cancer Center, Medical University of South Carolina (P30 CA138313). P.M. and S.S. were supported by NIH R21 CA286287-01A1. S.S. was supported by the American Cancer Society Institutional Research Grant: IRG-24-1290553-23-IRG. The content is solely the responsibility of the authors and does not necessarily represent the official views of the American Cancer Society, the National Cancer Institute, and the National Institutes of Health.

## Notes

### Competing Interest Statement

The authors have declared no competing interest.

